# Dendritic processing of spontaneous neuronal sequences for one-shot learning

**DOI:** 10.1101/165613

**Authors:** Tatsuya Haga, Tomoki Fukai

## Abstract

Spontaneous firing sequences colloquially called “preplay” are fundamental features of hippocampal network physiology. Preplay sequences have been hypothesized to participate in hippocampal learning and memory, but such functional roles and their potential cellular mechanisms remain unexplored. Here, we report a computational model based on the functional propagation of preplay sequences in the CA3 neuronal network. The model instantiates two synaptic pathways in CA3 neurons, one for proximal dendrite-somatic interactions to generate intrinsic preplay sequences and the other for distal dendritic processing of extrinsic sensory signals. The core dendritic computation is the maximization of matching between patterned activities in the two compartments through nonlinear spike generation. The model performs robust one-shot learning with long-term stability and independence that are modulated by the plasticity of dendrite-targeted inhibition. This model demonstrates that learning models combined with dendritic computations can enable preplay sequences to act as templates for rapid and stable memory formation.

## Introduction

Fast and robust memory encoding is a fundamental ability of the brain and has been extensively explored in the hippocampus. During spatial navigation, hippocampal place cells rapidly acquire their spatial receptive fields during the first exposure to the spatial environment (Nakazawa et al., 2003). In addition to this learning speed, hippocampus can form distinct spatial representations for multiple spatial experiences without interference (Alme et al., 2014; Mizuseki et al., 2012), avoiding overwriting of previous memories. Firing sequences exhibited by hippocampal place cells during locomotion (Foster and Wilson, 2007; Wang et al., 2014) are replayed in spontaneous activity (Carr et al., 2011; Lee and Wilson, 2002), and several studies have suggested the functional link between these firing sequences and the memory encoding process (Carr et al., 2011; Wang et al., 2014). Disrupting sharp wave ripples (Girardeau et al., 2009), which coincide with the replayed firing sequences, impairs memory consolidation, suggesting the importance of these sequences in memory formation.

Recent evidence has shown that a significant fraction of place-cell sequences emerge from firing sequences that are ‘pre-played’ in spontaneous activity prior to spatial experience (Dragoi and Tonegawa, 2011, 2013; Grosmark and Buzsáki, 2016; Ólafsdóttir et al., 2015). Although preplay sequences remain controversial (Silva et al., 2015), these results suggest that hippocampal networks may have an innate structure to utilize spontaneous firing patterns for sequence learning. Area CA3 is the likely source of prepay firing sequences in the hippocampus (Middleton and McHugh, 2016; Nakashiba et al., 2009; Omura et al., 2015), and CA3-based computational models have been proposed for sequence learning (Blum and Abbott, 1996; Bush et al., 2010; Gerstner and Abbott, 1997; Jahnke et al., 2015; Sato and Yamaguchi, 2003; Tsodyks and Sejnowski, 1995). However, these previous models studied simplified conditions in which the place fields of individual neurons were already configured prior to spatial exploration, and recurrent synapses learn sequential firing in an a posteriori manner without the involvement of spontaneous sequences. Therefore, while the functional role of replay sequences in hippocampal memory processing is relatively well understood (Carr et al., 2011; Girardeau et al., 2009), that of preplay sequences remains unknown. Indeed, the cellular mechanisms that would enable preplay-based memory processing has never been investigated.

In this study, we propose a computational model where hippocampal networks utilize preplay sequences for robust one-shot learning of spatial memory. Unlike previous models, our model does not assume that the place fields are preconfigured, but rather we hypothesize that spontaneous firing sequences exist prior to sequence learning, and the role of sequence learning is to associate sequences of sensory input patterns with specific preplay sequences. An unexpected finding of our framework is that dendritic computation between two segregated synaptic pathways to CA3 pyramidal cells is required to perform this pattern association. The perforant path from the entorhinal cortex conveys external sensory information to CA3 distal dendrites, while recurrent synapses primarily contact the CA3 proximal dendrites (Witter, 2007). This bipartite network architecture allows CA3 pyramidal cells to amplify and potentiate coincidence between two input streams, if CA3 pyramidal cells perform dendritic computation like calcium spikes in neocortical pyramidal cells (Larkum, 2013; Shai et al., 2015) and dendritic plateau potentials in hippocampal CA1 (Bittner et al., 2015; Takahashi and Magee, 2009). We construct a mathematically tractable and biologically plausible two-compartment neuron model by including Hebbian plasticity in each compartment (Oja, 1982) and implement canonical correlation analysis (CCA) (Hotelling, 1936; Izadinia et al., 2012) between the compartments, which emerges naturally from the effect of dendritic coincidence detection. The CCA-like learning maximizes matching between somatic spontaneous sequences and dendritic sensory inputs to rapidly form spatial memories, and increases the frequency of replay of the firing sequence associated with a particular experience. Dendritic inhibition stabilizes the place fields and protects distinct memory experiences from interferences. The model shows for the first time that a pivotal combination of dendritic computation and spontaneous firing sequences enable the rapid formation of stable place fields and place-cell sequences.

## Results

### Two-compartment neuron model with nonlinear dendritic computation

Dendritic coincidence detection and consequent synaptic plasticity play a pivotal role in the spatial memory encoding modeled below. Based on this principle, we constructed a mathematically tractable neuron model keeping its biological plausibility. We considered a two-compartment neuron model with a somatic compartment describing, in reality, the combination of a soma, basal and proximal dendrites, and a distal dendritic compartment representing the apical tuft dendrite. Assuming that the conductance between two compartments is small (Larkum, 2013), we modeled the activation of each compartment independently as

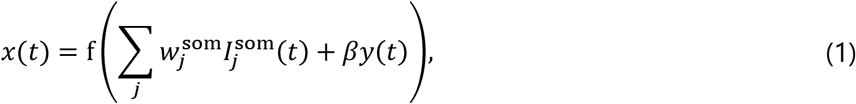

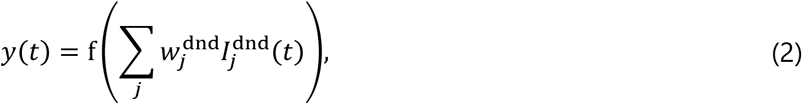

where *x(t)* determines the firing rate of sodium spikes in the somatic compartment, *y(ť)* is the local activity in the distal dendritic compartment, 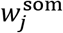 and 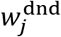 are synaptic weights on the somatic and dendritic compartments, respectively, and *βy(t)* is the threshold modification of somatic spikes by dendritic inputs(Shai et al., 2015). Unweighted postsynaptic currents 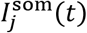 and 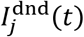 generated by each synapse are calculated by

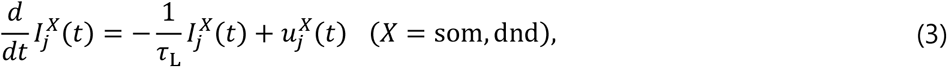

where τ_l_ = 10 ms is the decay constant of postsynaptic currents. Each neuron has a sigmoidal nonlinear response function *f(I)* = 1/(1 + exp(−(*I − θ_f_*))) with *θ_f_* being a constant threshold.

Synchronous activation of the two compartments represented by the product *x(t)y(t)* generates calcium spikes (or other similar dendritic mechanism for coincidence detection), which in turn enhance neuronal firing. Thus, the net output firing rate of the two-compartment neuron is expressed as

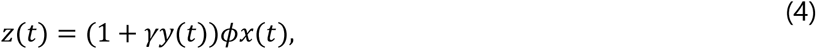

where *ɸ* is the maximum firing rate elicitable by local somatic inputs, and *γ* is the amplification factor of calcium spikes, and *z(t)* gives the output firing rate of the two-compartment neuron. We note that the above equation takes into account the experimental results that activation of distal dendrites increases the gain of the somatic firing rate (Larkum, 2013; Shai et al., 2015).

We express the learning rule for the two-compartment neuron as

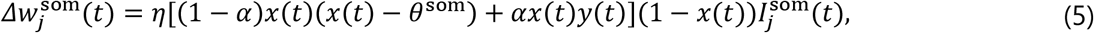

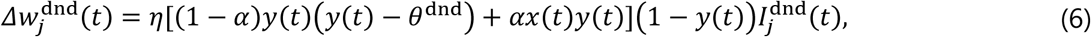

where *α* is a constant that determines the relative magnitude of the potentiation caused by calcium spikes. In both equations, the first terms in brackets represent Hebbian synaptic plasticity induced by local activities *x(t)* and *y(t)* by BCM theory (Bienenstock et al., 1982; Intrator and Cooper, 1992). Although BCM theory was originally introduced to describe the relationship between somatic firing rate and weight changes, calcium-based plasticity also gives a similar rule to BCM theory for dendritic activity (Yang et al., 2016). The second terms in brackets express the long-term potentiation (LTP) effect induced by calcium spikes generated by coincident inputs and successive high calcium ion influx into the dendrites. In hippocampal CA1 pyramidal neurons, the coincident activation of the perforant path projecting to apical tuft dendrites and the Schaffer collateral projecting to proximal dendrites induces LTP (Takahashi and Magee, 2009). Similar phenomenon was also observed in cortical neurons (Sjöström and Häusser, 2006).

Overall factors (*1 — x(t))* and (*1 — y(t)*), which do not change the direction of weight changes, were multiplied to match to the objective function we present later. As in the original BCM theory, moving thresholds 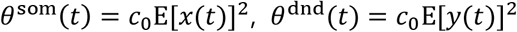 prevent run-away evolution of synaptic strength. We defined 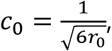, by which the mean activity in each compartment approximately converges to *r_0_* in the absence of coincidence detection, i.e., when *α* = 0 (see Methods). Homeostasis is also preserved for *α* > 0, although the mean activity becomes higher.

### PCA-like and CCA-like learning in the two-compartment neuron model

It is worth noting that the present learning rule for two-compartment neurons (*β* = 0) is derived from the following objective function:

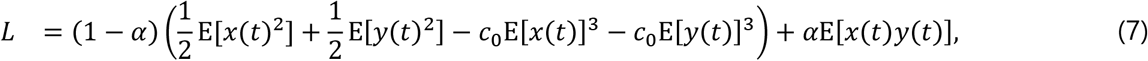

by gradient ascent

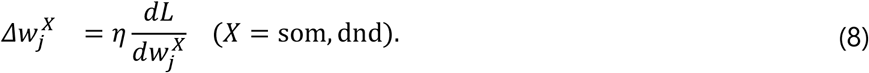

This objective function implies the maximization of second-order moments E[*x(t)*^2^], E[*y(t)*^2^] and correlation E[*x(t)y(t)*] in conjunction with the minimization of means E[*x(t)*]^3^ and E[*y(t)*]^3^. Therefore, the learning rule achieves the combination of PCA-like (Oja, 1982) and CCA-like (Hotelling, 1936; Izadinia et al., 2012) learning of input vectors 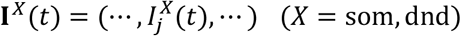 under a homeostatic constraint, where *α* determines the relative weight of CCA. In this paper, single-compartment neurons have only somatic compartment and hence perform only PCA-like learning supposed by BCM theory. In contrast, two-compartment neurons perform dual learning, that is, PCA-like learning within each compartment and CCA-like learning between the two compartments.

The learning behavior of the two-compartment neuron significantly varied depending on the correlation pattern of inputs. In Fig. 1a, the somatic and dendritic compartments received synaptic inputs from minority groups (A and A’) and majority groups (B and B’) of input neurons. Activities of these neurons were strongly correlated within each group but were uncorrelated between pairs of groups A-B’, B-A’ and A’-A’. We conducted simulations when A and A’ were either correlated or uncorrelated (Fig. 1b). When groups A and A’ were uncorrelated, synapses from groups B and B’ were strongly potentiated, but those from A and A’ were not (Fig. 1c, center). Accordingly, the activities of the two compartments were governed by inputs from groups B and B’, and hence were mutually uncorrelated (Fig. 1d, center). In this case, the learning performance was essentially the same as that of the single-compartment model (Fig. 1c, left: *α* = 0, no inter-compartment interaction). By contrast, when the activities of groups A and A’ were correlated, synapses from A and A’ were selectively potentiated whereas those from B and B’ were depressed (Fig. 1c, right). Accordingly, the two compartments exhibited correlated activities after learning (Fig. 1d, right). Note that in the two-compartment model output firing rate was approximately proportional to somatic activity.

**Figure 1:**
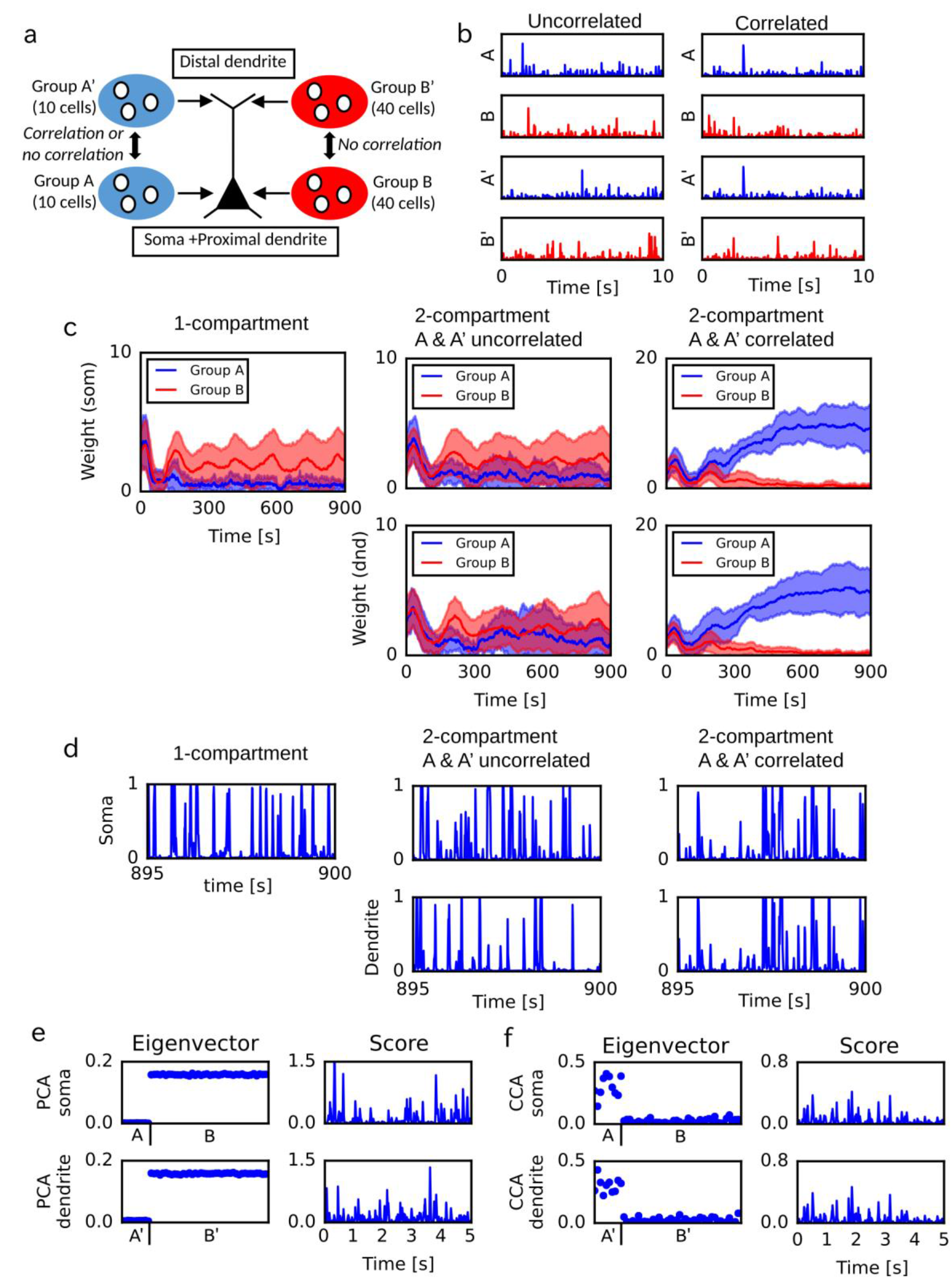
PCA-and CCA-like learning in a two-compartment neuron model. (a) Simulation settings. Each compartment receives inputs from 50 neurons. Each group A or A’ consists of 10 input neurons, and each group B or B’ of 40 neurons. (b) An example of input neuron activities when the groups A and A’ were correlated or uncorrelated. (c) Time evolution of synaptic weights are shown for the single-compartment model (left) and the two-compartmental model receiving uncorrelated (center) or correlated (right) inputs from A and A’. The single compartment neuron only received inputs from A and B. The means (lines) and standard deviations (shaded areas) are shown. (d) The activities of the neuron models were shown for the same simulation settings as in c. (e) PCA were applied to signals simulated in the same setting as in c when A and A’ were correlated. The eigenvectors (left) and the scores of the first PCs (right) are shown. (f) CCA were applied to signals simulated in the same setting as in c when A and A’ were correlated.

For comparison, we calculated the principal components of input vectors **I**^som^(*t*) and **I**^dnd^(*t*) when groups A and A’ were correlated. As expected, the first principal components extracted by PCA in the soma and dendrite were uncorrelated inputs from groups B and B’, respectively, and the scores (signals projected onto PC1 eigenvectors) were also uncorrelated between the two compartments (Fig. 1e). Then the pair of input vectors was analyzed by CCA, which extracted correlated inputs from groups A and A’ and also yielded highly correlated scores (Fig. 1f). Thus, CCA and the two-compartment neuron model operate similarly on correlated somatic and dendritic inputs.

These results imply that CCA-like learning of the two-compartment neuron model can extract a minor input component to one compartment if a coincident input is given to the other. The extraction of weak inputs based on correlation across compartments is a critical difference between our learning rule and conventional Hebbian learning, which basically extracts only major input components. However, if there is no coincident activity between the compartments, each compartment implements independent Hebbian learning and acts like an independent neural unit.

### The role of inhibitory feedback in the two-compartment neuron model

Hippocampus has interneurons serving perisomatic and dendritic inhibitory feedback projections (Müller and Remy, 2014; Royer et al., 2012). We modeled these inhibitory feedback mechanisms in the two-compartmental neuron model (Fig. 2a) to realize the stability and functional specialization of dendritic synapses. In reality, interneurons targeting the soma and dendrites of pyramidal neurons belong to different cell types. For simplicity, however, we only consider a single population of inhibitory neurons in our network model. We approximated the net output from this population *I*^inh^(*t*) by the sum of the output from all pyramidal neurons. Pyramidal neuron *i* was modeled as a two-compartment model with inhibitory feedback:

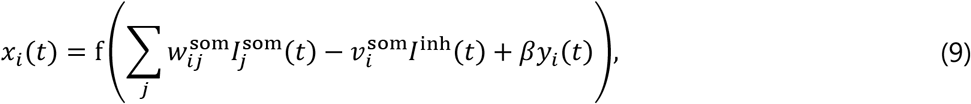

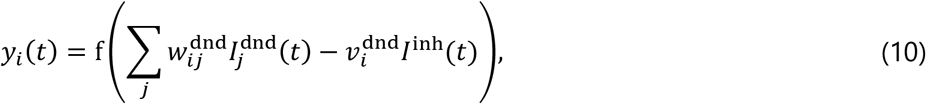

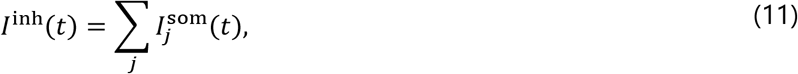

where 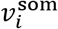 and 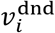 are inhibitory synaptic weights.

**Figure 2:**
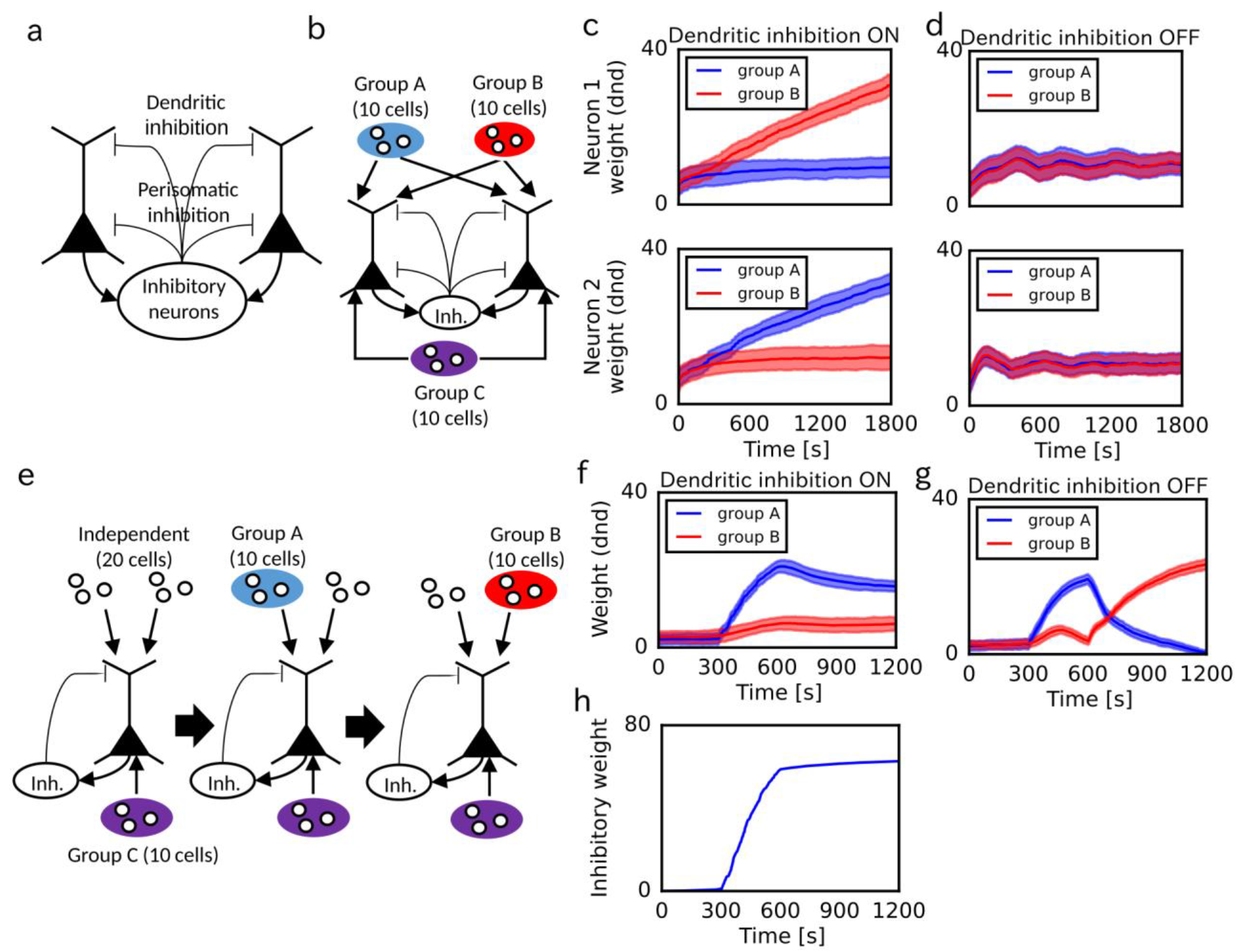
Roles of plastic inhibitory feedback to the distal dendritic compartment. (a) Inhibitory feedback model is schematically illustrated. Inhibitory interneurons project to both somatic and dendritic compartments. (b) In this simulation setting, two pyramidal neurons projected to an inhibitory neuron and received inhibitory feedback at the somatic and dendritic compartments. In addition, pyramidal neurons received common somatic inputs from excitatory cell group C and mixed dendritic inputs from two mutually-uncorrelated excitatory cell groups A and B. The activity of cell group C was correlated with the activities of cell groups A and B with equal magnitudes. (c, d) Time evolution of synaptic weights on the dendritic compartments of the two cells with (c) or without (d) dendritic inhibition. The means (lines) and standard deviations (shaded areas) of synaptic weights are shown. (e) A single pyramidal neuron with inhibition fed back onto its dendrite received somatic inputs from a cell group C and dendritic inputs from two cell groups A and B. Activities of input neurons in groups A and B were initially uncorrelated within each group and with other groups. At time 300 sec, correlations were introduced within group A and between groups A and C. At time 600 sec, neurons in group A returned to an uncorrelated state, but neurons in group B became correlated within the group and with group C. (f, g) Time evolution of excitatory synaptic weights on the dendritic compartment with (f) or without (g) dendritic inhibition. (h) Time evolution of inhibitory synaptic weights on the dendritic compartments is displayed.

It has been observed experimentally that not only excitatory but also inhibitory synapses exhibit activity-dependent plasticity (Vogels et al., 2013). Although the property of inhibitory synaptic plasticity has not been fully understood, here we assumed that inhibitory weights for the distal dendritic compartment are modified by a similar learning rule to excitatory synapses:

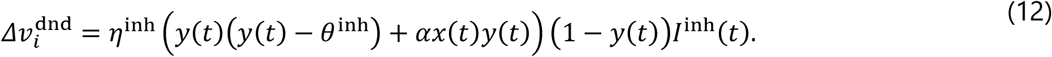

In this expression, θ^inh^ is a constant threshold and was fixed at 0.5 throughout this research (we note that the choice of this parameter value is not crucial for the performance of this model). We did not implement the plasticity for perisomatic inhibitory synapses to avoid the instability of network’s learning behavior.

To clarify the role of the plastic dendritic inhibition in CCA-like learning, we tested the behavior of the neuron model in two conditions. First, we simulated a coupled network of two pyramidal neurons and a population of inhibitory neurons (Fig. 2b). When the somatic input was correlated with both dendritic inputs with equal magnitudes, these neurons selectively learned different dendritic inputs in the presence of dendritic inhibition (Fig. 2c), but not in its absence (Fig. 2d). At a first glance, the result is a mere extension of feature extraction by lateral inhibition in competitive networks of single-compartment neurons. Actually, the present model had lateral inhibition between the soma and between the dendrites, and somatic activity was amplified by the activation of dendrites (Eq. 4). Nevertheless, here the intersomatic lateral inhibition alone was unable to separate the dendritic inputs in CCA-like learning. Thus, dendritic inhibition is required for the functional specialization of dendrites.

In the second case, we examined the robustness of dendritic excitatory synapses against changes in correlation structure of synaptic inputs (Fig. 2e). Without dendritic inhibition, an abrupt change in correlations between somatic and dendritic inputs triggered a significant reorganization of dendritic excitatory synapses (Fig. 2g). By contrast, such a reorganization did not occur in the presence of dendritic inhibition (Fig. 2f). This stability was due to the potentiation of dendritic inhibitory synapses during the learning of the initial correlation structure between groups A and C (Fig. 2h). The potentiated dendritic inhibition changed the excitation-inhibition balance of the dendritic compartment such that its responses to a learned input pattern were enhanced whereas those to other input patterns were suppressed.

Thus, in our model the potentiation of dendritic inhibition separates and stabilizes receptive fields on the dendrites acquired by CCA-like learning. We will show later that these properties play crucial roles in the robust memory encoding in a recurrent network model of the two-compartment neurons.

### Robust one-shot learning of place fields by two-compartment neural network

Using the neuron model and the inhibitory feedback model described above, we constructed a CA3 recurrent network model to investigate the role of our two-compartment model in sequence memory (Fig. 3a). In this model, the somatic compartments of pyramidal neurons receive excitatory recurrent connections, input from the dentate gyrus (DG), theta-band (7 Hz) oscillatory input from the medium septum (Wang et al., 2014) and random noise, while the dendritic compartments receive input from the entorhinal cortex (EC). Excitatory connections are reciprocally wired such that the recurrent network can propagate firing sequences (Romani and Tsodyks, 2015; Wang et al., 2014). In the presence of noise, Poisson firing of DG activates a small portion of CA3 neurons, which triggers preexisting firing sequences without EC input (Fig. 3b). During locomotion, DG input conveys contextual-dependent information in the sense that DG is only activated at particular spatial positions (see below). Firing sequences are modulated by theta-band oscillations due to the medial septal input. During the propagation, we decreased the release probability of neurotransmitters at recurrent connections to suppress the decay speed of synaptic weights. In the hippocampus, acetylcholine is known to modulate glutamatergic neurotransmitter release (Bush et al., 2010; Hasselmo, 2006).

**Figure 3:**
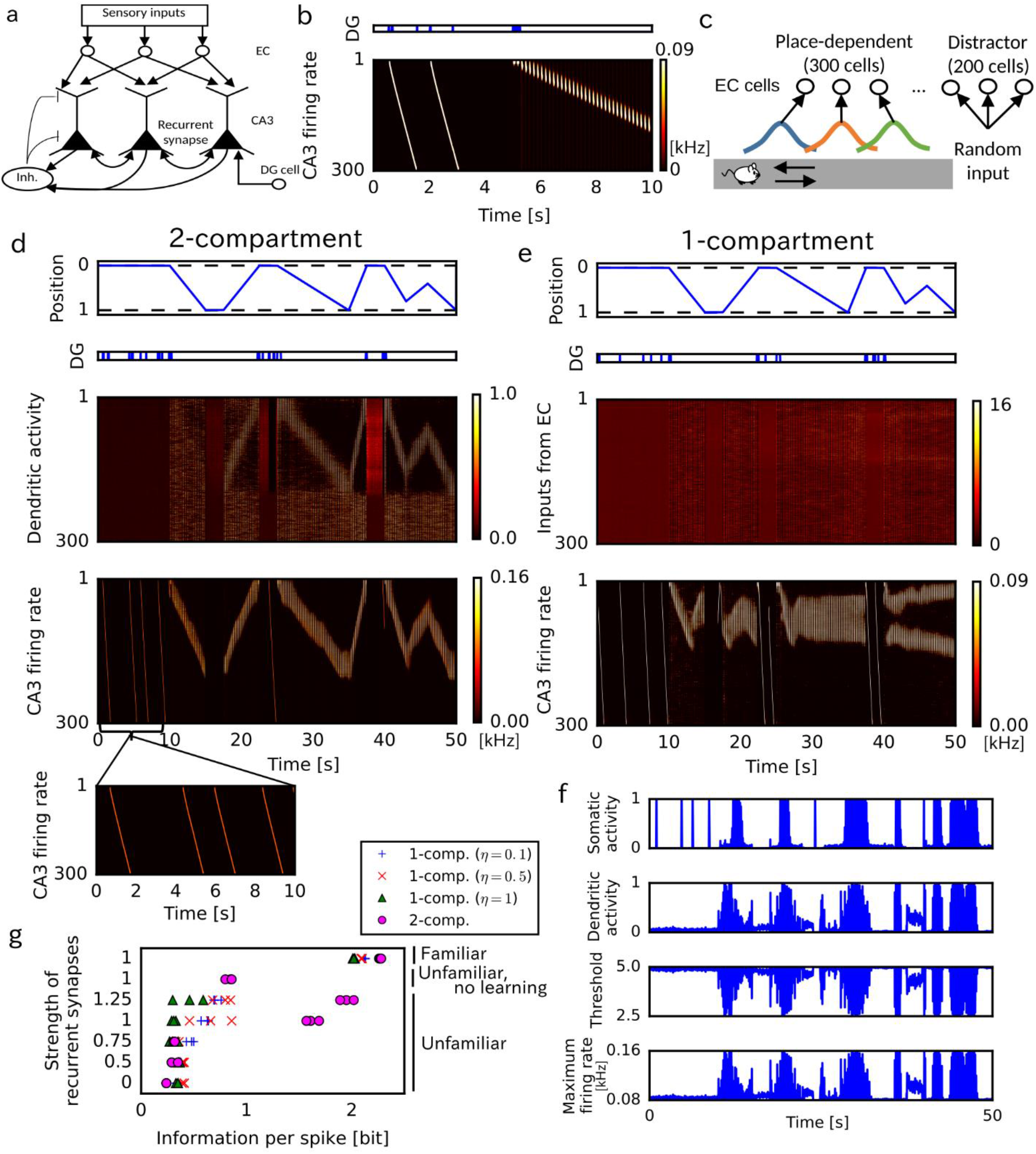
One-shot learning of place fields on a one-dimensional track. (a) Our CA3 network model consists of 500 EC neurons projecting to the distal compartments of 300 two-compartment CA3 neurons, which have inhibitory feedback to both distal dendritic and somatic compartments. DG input activates neuron 1 to neuron 10 of CA3 in a probabilistic manner. (b) DG-evoked preexisting activity patterns in CA3 were simulated without EC input. The animal was immobile from 0 to 5 sec and ran from 5 to 10 sec. (c) The behavioral paradigm and activities of EC neurons in the present simulations. Position-dependent sensory features are encoded by 300 EC neurons, whereas other 200 EC neurons (neuron ID 300 to 500) show position-independent distractor activity. (d,e) Activities of the two-compartment network model (d) and single-compartment network model (e) for animal’s movements shown in the top panels. The single-compartment network model was simulated with *η* = 0.5. The inset in (d) shows the expanded plot of the output firing rate from 0 s to 10 s. (f) Time evolution during learning is shown for the dynamical variables of the two-compartment neuron. The examples were from CA3 neuron #100. (g) Average information per spike was calculated in various conditions. Three simulation trials were performed in each condition with different initial conditions. The strength of recurrent connections was measured relative to the connection strength used in c and d. In simulating familiar tracks, we used the initial weights of EC-to-CA3 synapses optimized to generate place-dependent firing. In the simulations of unfamiliar tracks, these initial weights were randomly shuffled.

We considered an animal running back and forth on a one-dimensional (1D) track. During a run, some EC neurons are sequentially activated by position-dependent sensory features on the track, while others (distractors) are activated randomly (Fig. 3c). Prior to the first run, there is no way for the animal to know the sensory features and their order of appearance along an unfamiliar track. Therefore, the initial weights of EC-to-CA3 projections were chosen randomly, and accordingly dendritic activity showed no place-dependence initially. The position-dependent EC activity may correspond to the representation of local landmarks in the lateral entorhinal cortex or the firing fields of grid cells in the medial entorhinal cortex (Knierim et al., 2014).

Initially, the animal was immobile at an endpoint of an unfamiliar track, where DG exhibited a brief and repeated activity (Fig. 3d, top), which in turn activated the preexisting sequences in spontaneous CA3 activity (Fig. 3d, bottom). These sequences were initially not associated to any sensory information (hence any spatial information) represented in EC. However, during the first traversal the dendritic compartments learned to associate sequential sensory inputs from EC with the firing sequence triggered by DG (Fig. 3d, middle). During subsequent runs, the activity of distal dendrites established control of somatic activity, and hence of sequence propagation, by modulating the gain and threshold of somatic firing rates (Fig. 3f). Consequently, CA3 neurons showed clear place-dependent firing in the second and third traversals even though this was the first exposure to an unfamiliar track and synapses from EC were initially random and not pre-configured to any convenient spatial pattern. We note that the reorganization of EC-to-CA3 projections enabled firing sequences to follow abrupt changes in running speed and directions in the second and third traversal.

For comparison, we constructed a single-compartment network model and trained it on the same spatial navigation task. In this model, both recurrent synapses and EC inputs were connected to the somatic compartments (thus, the dendritic compartments were passively driven by the somatic compartments and did not play any active role). This model failed to form place fields in the present simulations (Fig. 3e). For the relatively large value of the learning coefficient used in the simulations, the formation of place fields was easily disturbed by noise (from the distractor EC neurons) before they became robust. On the other hand, if the learning coefficient was small, firing sequences could not follow changes in the movement directions of the animal at both ends of the maze.

We assessed the quality of the place fields formed in various simulation conditions by means of “information per spike” (Skaggs et al., 1993), a measure based on the mutual information between neural activity of each cell and animal’s position (see Methods). As shown in Fig. 3g, in both models the average mutual information was high in a familiar track, which we simulate the CA3 activity with optimized initial synaptic weights of EC-to-CA3 projections. In an unfamiliar track, however, only the two-compartment model acquired highly place-dependent neural activity as in the familiar track, whereas the single-compartment model exhibited low mutual information for all three values of the learning coefficient. Learning of the unfamiliar track was required for the place dependence as the performance of the two-compartment model was impaired by eliminating the plasticity effect *(η* = 0). Importantly, when we decreased the initial weights of recurrent synapses from the original setting (by a multiplicative factor of 0.75, 0.5 or 0), the two-compartment model also failed to acquire place-dependent activity on the unfamiliar track. On the other hand, increasing the weights (1.25 times) did not degrade the performance in learning. These results imply that the activity patterns preexisting in the recurrent network are crucial for the place field formation in the two-compartment model. Conversely, dendritic computation is necessary for the efficient use of preexisting sequences for memory formation.

### Long-term stability of memory in remote replay events

At a first glance, one-shot learning looks easily achievable by sufficiently fast synaptic modifications. However, this was not the case as there was a trade-off between learning speed and the long-term stability of memory. In the case of spatial memory, the place fields formed in a previous awake state have to be preserved during spontaneous replays in sleep states (Lee and Wilson, 2002) and awake replays of remote experiences (Carr et al., 2011). In this section, we examine the stability of spatial memory against such replay events. To this end, we trained the network model on one-shot encoding of an unfamiliar track (Fig. 4a). We then introduced random noise in EC neurons and the CA3 network to generate irregular activity in EC and spontaneous firing sequences in CA3 (Fig. 4b). After exposing the network to random inputs and spontaneous replay events for 600 sec, we tested it for the exploration of the learned track.

**Figure 4:**
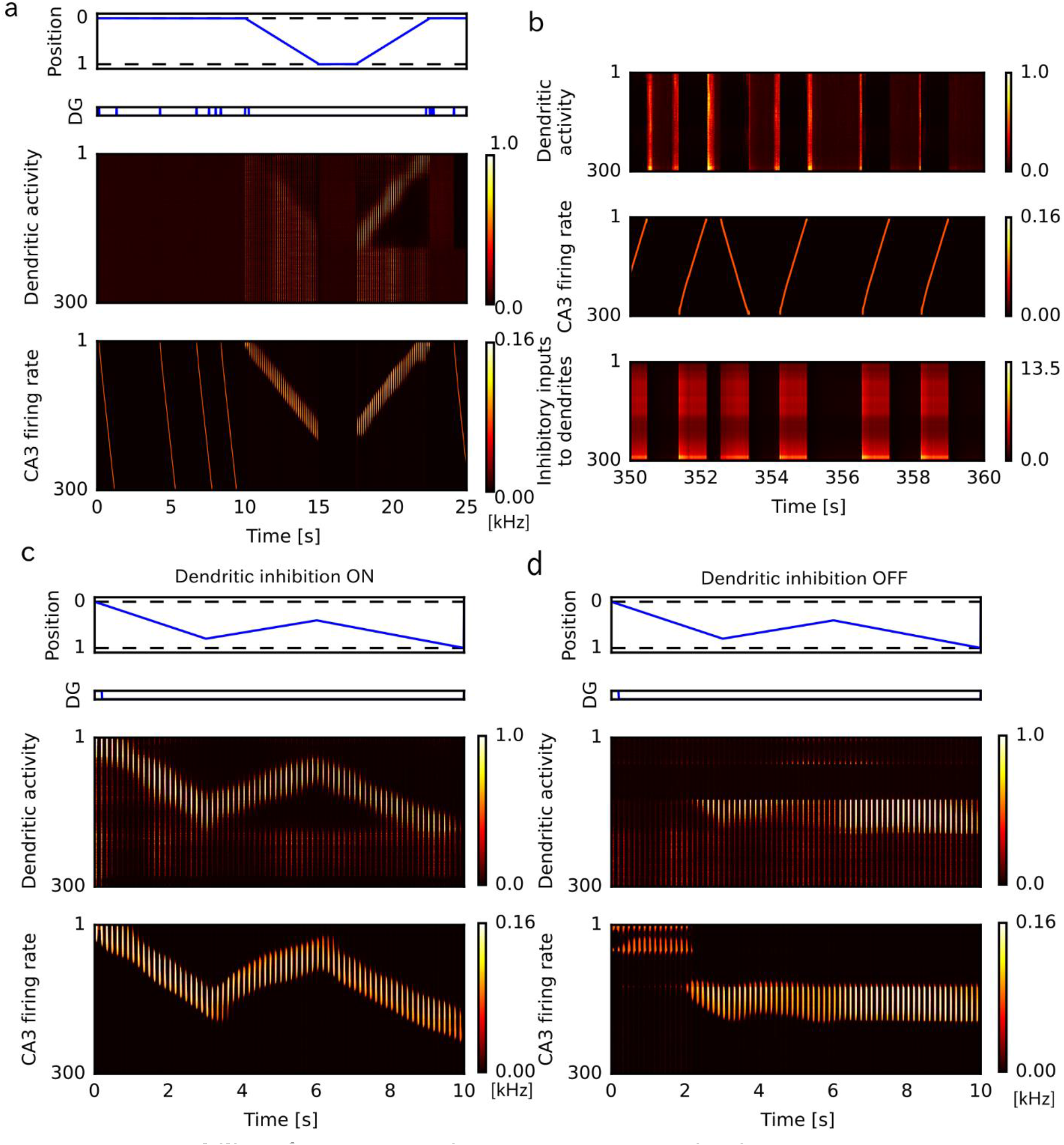
Long-term stability of memory against spontaneous activation. (a) Dendritic and somatic activities of the two-compartment CA3 neurons are shown before and during the first traversal on a one-dimensional track. (b) Dendritic activity and firing sequences during spontaneous activity are shown together with inhibitory inputs to the dendritic compartments. (c) Activity of the two-compartment network model during traversals on the one-dimensional track is shown after exposure to spontaneous activity. (d) Similarly, such network activity is shown in the case that the dendritic inhibition was removed during the exposure to spontaneous activity.

In spite of the repetitive coupling of random inputs and replay events, the two-compartment neurons still preserved their place fields after exposure to random noise (Fig. 4c). This was achieved by the suppression of dendritic activity during replay events by lateral inhibition between the dendritic compartments (Fig. 4b, bottom), as we showed in Fig. 2. The inhibitory effect prevented an undesirable association of random EC inputs and spontaneous replays in CA3. Actually, the place fields were completely eliminated when all inhibitory weights were set equal to zero during replay events (Fig. 4d). Thus, the results of this and previous sections demonstrate that our two-compartment network model reconciles conflicting demands on the brain’s memory systems, i.e., one-shot learning and the long-term stability of memory, without an ad hoc tuning of model parameters.

### Plasticity in dendritic inhibition prevents overwriting of multiple episodes

So far, we have studied the one-to-one association of a linear track and a firing sequence. However, in many real-world tasks, the hippocampus has to separately store multiple memories. In spatial navigation experiments, CA3 has been shown to develop sparse and orthogonal spatial representations (Alme et al., 2014; Mizuseki et al., 2012). To see whether our two-compartment neuronal network is capable of learning such representation in complex spatial environments, we tested the formation of spatial memory in the case where the animal visits three arms on a Y-maze in turn and repeatedly (Fig. 5a). We configured initial recurrent connections such that CA3 network had three preexisting firing sequences (Fig. 5b), which is a minimal requirement for learning the branching spatial structure. DG was assumed to fire at the junction of the Y-maze, and DG input at the center could trigger all these firing sequences simultaneously. The preexisting sequences were spontaneously generated by noise and DG inputs in the resting state (Fig. 5c), and the firing sequences were accompanied by theta oscillation during running back and forth on each arm (Fig. 5d-f), as in the previous simulations of 1D maze. We note that intersomatic lateral inhibition prevented the coactivation of all three preexisting sequences by input from DG, and the choice from three sequences was initially determined by random noise.

**Figure 5:**
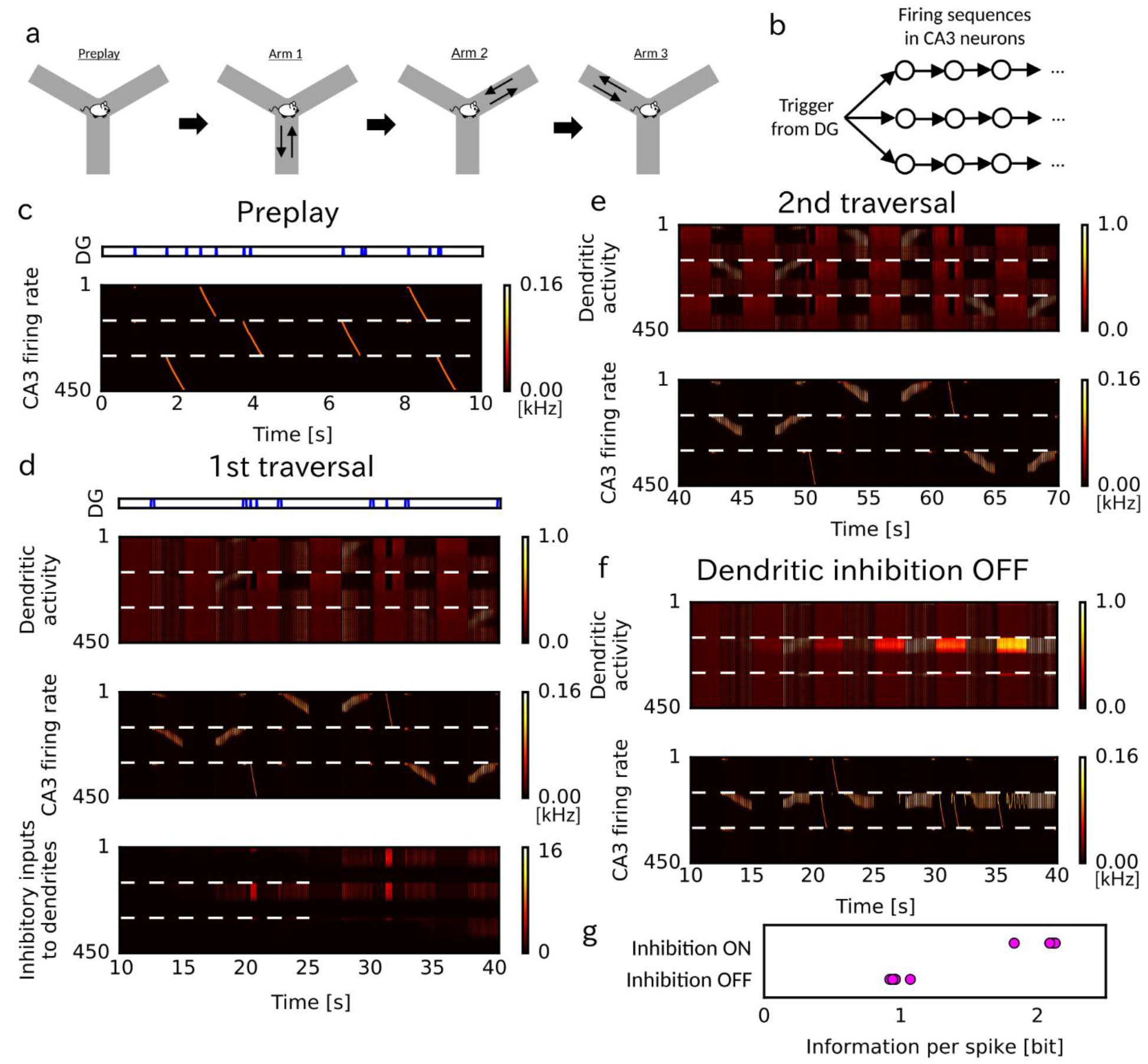
Orthogonal memory formation in a Y-maze. (a) Behavioral paradigm is schematically illustrated. The animal starting from the junction successively visits three arms on a Y-maze. (b) In the initial setting of the recurrent network, a DG input triggers three firing sequences when the animal is at the junction of the Y-maze. Each branch of sequence consists of 150 neurons. (c) Spontaneous activity is shown for the three branches of the two-compartment network model before the exploration. (d) The dendritic and somatic activities of the two-compartment network model during the first run on the Y-maze. Time evolution of inhibitory inputs to the dendritic compartments is also shown. (e) Activity of the two-compartment network model during the second run is shown. (f) Activity of two-compartment model without dendritic inhibition during the first run on the Y-maze. (g) Information per spike was calculated over five simulation trials with and without dendritic inhibition.

The two-compartment network model robustly assigned the individual firing sequences to representing different arms (Fig. 5d and 5e). Inhibitory plasticity played a crucial role in the learning procedure. After the first traversal on an arm, a firing sequence or an ensemble of neurons is assigned to the arm. Then, dendritic inhibition decreases the response gain of these neurons in other arms (Fig. 5d, bottom) and selects such firing sequences that have not been associated with any spatial experience for learning the second arm. Because the trigger from DG to the soma is shared, separate learning of dendritic input patterns to different neurons requires dendritic inhibition, as we showed in Fig. 2. Thus, as shown in Fig. 5f, the formation of independent memory representations (firing sequences) for different spatial experiences is not guaranteed without dendritic inhibition. We performed five simulations with and without dendritic inhibition using different random seeds and always obtained qualitatively the same results (in Fig. 5g, the learning performance was assessed by information per spike). By this mechanism, every time the animal encounters a novel spatial experience, this network model exhaustively encodes it into a yet unassigned firing sequence, avoiding to overwrite old episodes with a novel episode.

### Replay of firing sequences is biased by recent experiences

Does our network model change spontaneous activity patterns in an experience-dependent fashion consistent with experiment? The structure of correlations in spontaneous hippocampal activity is different before and after experiences (Wilson and McNaughton, 1994). In particular, replay sequences become statistically significant only after experiences (Silva et al., 2015). Because the memory encoding of our model strongly relies on preexisting firing sequences, we examined whether our model is consistent with these findings in the case that that the CA3 recurrent network has somewhat complex structure. To be more specific, we considered a two-compartment network model with bifurcating firing sequences (Fig. 6a). A neuron at the junction was initially connected to both bifurcating pathways with equal strength, propagating firing sequences into one of the branches with approximately equal probabilities (Fig. 6b). After the animal explored the 1D track, the model associated one of the branching sequences with this experience (Fig. 6c, input pattern 1). Spontaneous activity selectively replayed this firing sequence after this experience (Fig. 6d), implying that the recurrent connections responsible for this sequence were selectively potentiated during the episode. We simulated the network model for five different sets of firing sequences, and observed strongly selective replay of the associated sequences (Fig. 6f). The maximum weight change in recurrent connections was 10 to 20 % of the maximum initial weight (data not shown), which was not large enough to create novel firing sequences but was large enough to modulate the probability of sequence propagation into different branches.

**Figure 6:**
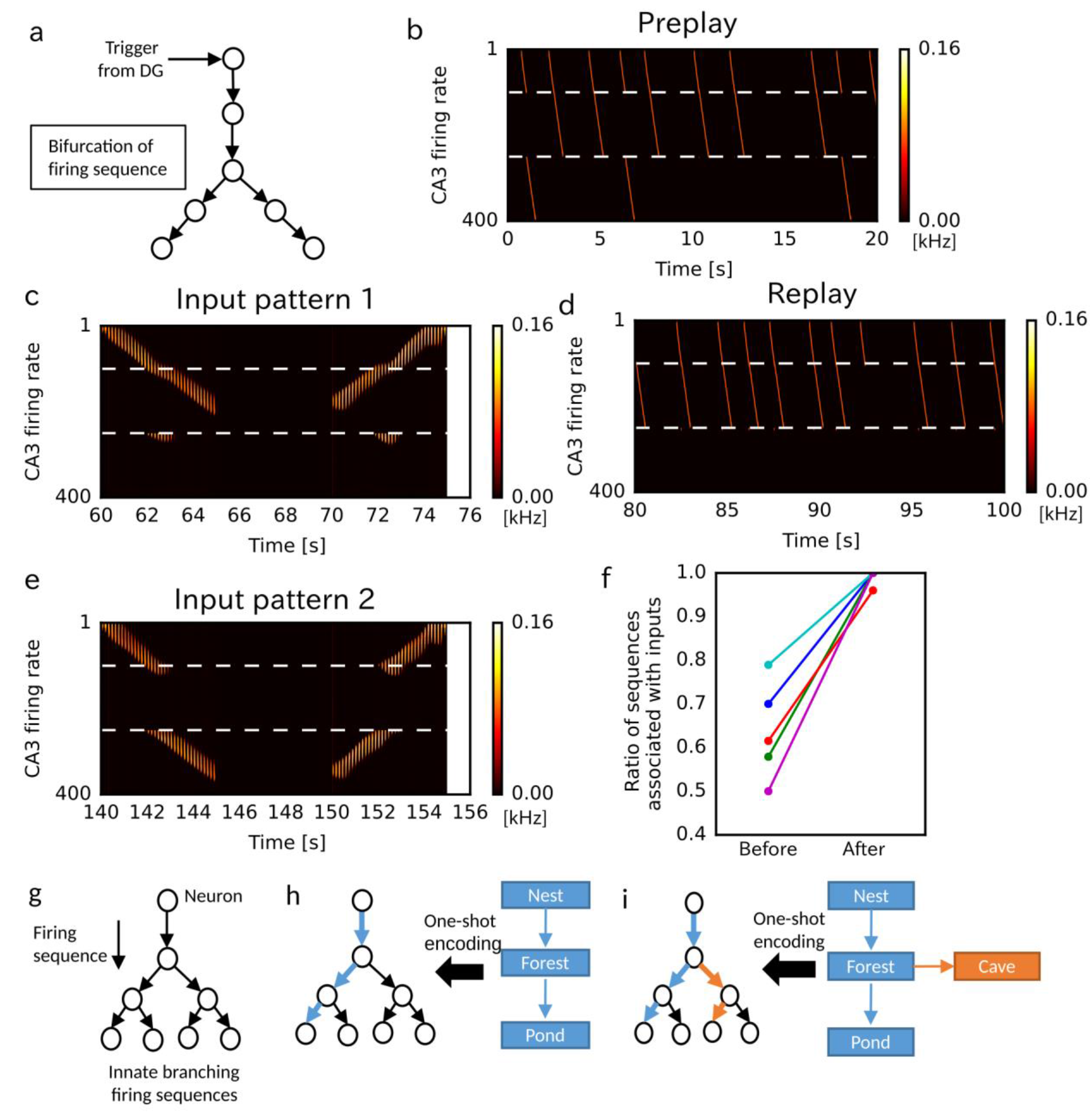
Memory encoding on branching firing sequences. (a) The somatic recurrent network of the two-compartment neuron model that has a bifurcating point. Neurons 1 to 100 constitute the trunk, 101 to 250 the left-side branch, and 251 to 400 the right-side branch. (b) Spontaneous branching firing sequences before spatial exploration are shown. (c) The two-compartment network model associated a sequence of sensory events (input pattern 1) is shown with the left-side branch of synaptic pathways. (d) After this encoding, spontaneous replay was biased to the firing sequence associated with input pattern 1. (e) The network model encoded a novel sensory sequence (input pattern 2) into the right-side branch of synaptic pathways. (f) The relative frequency of replay of the spontaneous firing sequence encoding input pattern 1 was calculated before and after the first experience for five simulation trials using different random seeds. The numbers of sequences propagating into either branch were counted for 60 sec in spontaneous activity. (g) The proposed memory encoding model utilizes a rich repertoire of branching firing sequences in the CA3 network. (h) Sequential sensory events are associated to a branching of firing sequences. (i) Novel sensory events are encoded into a different branch of sequences.

Next, we investigated how the model trained on input pattern 1 represents novel changes in the environment. To this end, we randomly shuffled the temporal order of sensory objects appearing along the 1D track (equivalently, the temporal order of firing in EC neurons). Our model encoded the novel sensory sequence (input pattern 2) into another branch that had not been previously used (Fig. 6e).

Assuming that the innate recurrent network of CA3 is capable of generating a rich repertoire of firing sequences (Fig. 6g: see the discussion on this point), we propose the following memory encoding mechanism for hippocampal circuits. When an animal experiences sequential sensory events (for instance, from the nest to a pond through a forest), these events are rapidly associated with a branching firing sequence that happens to be most strongly correlated with the events (Fig. 6h). After this rapid encoding, plasticity at recurrent connections reorganizes the network structure to spontaneously replay this sequence more often than others, which further increases the robustness of the sequence, keeping receptive fields by dendritic inhibition. Whenever the animal experiences the same sensory sequence, the particular firing sequence associated with this sensory experience is reactivated. Now, consider the case where the animal chooses another path at some spatial position (for instance, from the nest to a cave through the forest). Our model suggests that sensory events on the novel path are assigned to a different branch of firing sequences (Fig. 6i). This model has several merits. First, memory encoding is fast and robust, as demonstrated in this study. Second, no additional neural resources are required for encoding the experienced part of novel experiences (i.e., from the forest to the cave). Third, during the update of spatial map, the spatial relationships between old and novel sensory objects are naturally preserved in the existing branching structure of neural networks. Thus, the concept of “preplay” reconciles with “experience-dependent replay” in the proposed mechanism.

## Discussion

In this study, we propose a novel framework of hippocampal memory processing based on a two-compartment neuron model that incorporates the effects of dendritic spikes into Hebbian learning. Our model of dendritic computation gives a plasticity rule that combines canonical correlation analysis of correlated somatic and dendritic inputs with the conventional Hebbian plasticity rule for PCA of uncorrelated inputs. A recurrent network model of the two-compartment neurons demonstrates robust one-shot learning of sequential sensory events by utilizing spontaneous firing sequences (i.e., preplay events). The model predicts that inhibitory plasticity at the dendrites of pyramidal cells plays pivotal roles for the stability and functional specialization of dendritic activity during learning. Our results indicate that dendritic computation is a crucial element of fast and robust memory encoding into preexisting hippocampal circuits.

### Mechanisms and functional implications of CCA-like learning

In the two-compartment neuron, CCA-like learning extracts a minor input component at one compartment when correlated input is given to the other compartment. We built this model based on experimental studies in neocortex (Larkum, 2013; Shai et al., 2015; Sjöström and Häusser, 2006) and hippocampal CA1 (Bittner et al., 2015; Takahashi and Magee, 2009). In cortical pyramidal neurons, weak synaptic inputs to the apical tuft are strongly attenuated in the apical shaft and usually cannot generate somatic sodium spikes. However, calcium spikes initiated in a “hot spot” in the apical trunk reliably propagate to the soma to generate a burst of sodium spikes (Larkum, 2013). Although the generation of calcium spikes usually requires a strong activation in the apical tuft, the threshold of calcium spikes decreases during the 30-ms time window around somatic sodium spikes, allowing weak inputs onto apical tuft to initiate calcium spikes (Larkum, 2013). In hippocampal CA1, coincident inputs from CA3 and EC initiates dendritic plateau potentials which also drive burst firing (Bittner et al., 2015). These dendritic computation can detect and amplify coincident activation of soma and distal dendrites. Furthermore, dendritic coincidence detection induces LTP in both cortex (Sjöström and Häusser, 2006) and hippocampus (Takahashi and Magee, 2009). Such dendritic mechanism for coincidence detection is also essential for the rapid formation of place fields in our CA3 network model. The proposed role of dendritic computation in the rapid formation of place fields was recently supported in CA1, where artificially induced dendritic spikes generated arbitrary place fields in pyramidal neurons (Bittner et al., 2015).

Similar learning schemes are also expected to work in the neocortex. Calcium spikes presumably contribute to the integration of functionally distinct inputs to the basal dendrites and apical tufts (Larkum, 2013). In engineering, CCA is a well-established multivariate analysis method used in variety of applications such as the integration of multi-modal sensory inputs in video streams (Izadinia et al., 2012). Our two-compartment model suggests that CCA is also important in the brain and proposes a cellular mechanism of CCA based on the physiology of pyramidal-cell dendrites.

### Crucial roles of CA3 dendritic inhibition in sequence memory encoding

We suggest that activity-dependent inhibitory plasticity at the distal dendrites of pyramidal cells prevents the overwriting of memories, which is a long-standing issue in memory processing (Benna and Fusi, 2016). In our model, the plasticity of dendritic inhibition is crucial for the stability of memory traces (Fig. 4). Moreover, dendritic inhibition is important to robustly associate multiple sequential experiences with multiple firing sequences of different neuron ensembles without interferences or overwriting (Fig. 5). Somatostatin-positive (SOM+) interneurons target the apical dendrites of pyramidal cells in the hippocampus (Müller and Remy, 2014; Royer et al., 2012). This interneuron subtype is likely to provide the dendritic inhibition.

The dendritic computation modeled here may revise the conventional view of hippocampal microcircuit function. Area CA3, with rich recurrent connections, is generally referred to as a pattern completion system (Guzman et al., 2016; Nakazawa et al., 2002) in contrast to the DG, which is thought to be a pattern separation system (McHugh et al., 2007). However, our model suggests that the CA3 network can also perform pattern separation because, with the plasticity of dendritic inhibition, different memory items can be allocated to different CA3 neurons in spite of the same trigger input from DG. In our model, DG contributes to context-dependent separation and offers a trigger to CA3, where sensory inputs from the EC are further separated within the same context by dendritic computation. Thus, pattern separation is performed at multiple stages of hippocampal memory processing. Joint contributions of DG and CA3 to pattern separation was also suggested in a recent experiment (Senzai and Buzsáki, 2017).

### The network mechanism for preplay sequences

Although the relation between firing patterns of place cells during run and preexisting firing sequences has been shown (Dragoi and Tonegawa, 2011, 2013; Grosmark and Buzsáki, 2016; Ólafsdóttir et al., 2015), some other studies suggest firing patterns are organized through spatial experiences (Silva et al., 2015; Wilson and McNaughton, 1994). In modeling studies, the CA3 recurrent network with log-normal synaptic weight distributions can generate tremendously many spontaneous firing sequences with various branching patterns (Ikegaya et al., 2013; Omura et al., 2015; Teramae et al., 2012), and experimental evidence suggests such a distribution in CA3 (Ikegaya et al., 2013). Therefore, the innate structure of the CA3 recurrent network is expected to generate spontaneous firing sequences prior to learning, and our model suggest that encoding of sequential experiences into these preplay sequences naturally occur under the assumption of dendritic coincidence detection. Furthermore, we demonstrated that even the one-time experience can change the bias in branching firing sequences without major rewiring of the network structure. If spontaneous activity of naïve CA3 has the complex branching structure, this bias towards the branch used for memory may explain the change of correlation structure between hippocampal neurons. Although temporal coding in CA1 depends on inputs from CA3 (Middleton and McHugh, 2016), it is also possible that plasticity of CA1 synapses modulates the firing patterns.

It will be intriguing to study whether realistic log-normal networks can provide a repertoire of firing sequences that is sufficient for memorizing complex sensory experiences. Such recurrent network will also allow us to test whether our model generates place fields in 2D environments. Previous models (Brunel and Trullier, 1998; Káli and Dayan, 2000) have shown that Hebbian plasticity reorganize somatic recurrent connections to generate omnidirectional 2D place fields from multiple 1D place fields passing through a particular position from different angles. The same mechanism is expected to work in our model if recurrent network has a rich repertoire of firing sequences to learn pathways in different angles, although the formation of stable place fields will be much slower than learning 1D tracks. This reorganization may result in recruitment of new cells to firing sequences through learning (Grosmark and Buzsáki, 2016). Simulations of such network models, however, require a spiking version of the two-compartment model and an efficient platform for large-scale network simulations.

### Testable assumptions and predictions

The most important assumption of our model is the dendritic mechanism for correlation maximization (CCA), which was modeled based on findings in neocortex and CA1. Although there are some related experimental studies in CA3 (Kim et al., 2012; Makara and Magee, 2013), whether dendritic computation in CA3 pyramidal cells is analogous to that in CA1 and neocortical pyramidal cells should be clarified by future experiments.

Our model also assumes the potentiation of both excitatory and inhibitory synapses by coincident somatic and dendritic activation. While inhibitory plasticity depends on calcium signals (Kurotani et al., 2008; Sieber et al., 2013), whether it depends on dendritic spikes has yet to be examined in the hippocampus. Our results predict that the loss of dendritic inhibition disrupts the stability and orthogonality of CA3 place fields. Whether the removal of dendritic inhibition in CA3 triggers forgetting or remapping of spatial memory before consolidation is an interesting open question. Selective deletion (Cichon and Gan, 2015) or optogenetic inactivation(Royer et al., 2012) of SOM+ interneurons may remove dendritic inhibition. Alternatively, activation of vasoactive intestinal polypeptide-positive (VIP) interneurons which disinhibit distal dendrites (Yang et al., 2016) may lead to similar results.

Our model suggests that plasticity of EC-to-CA3 synapses is more important than that of recurrent synapses in CA3 for one-shot learning of place fields. Though the ablation of NMDA receptors in CA3 are known to result in the disruption of pattern completion and one-shot learning (Nakazawa et al., 2002, 2003), which connections are more responsible, recurrent synapses or EC-to-CA3 synapses, for one-shot learning has to be yet clarified.

### Relationship to other models of CA3 and dendritic computation

Previous models learn sequences under the assumption that place fields are configured prior to learning (Blum and Abbott, 1996; Bush et al., 2010; Gerstner and Abbott, 1997; Jahnke et al., 2015; Sato and Yamaguchi, 2003). In unfamiliar environments, this assumption is valid after grid cells, which estimate the self-position by path integration (Knierim et al., 2014), are formed in the medial EC. However, grid cells mature slower than place cells (Langston et al., 2010; Wills et al., 2010) and grid firing patterns requires an excitatory drive from place cells (Bonnevie et al., 2013). Our model demonstrated that place cells are rapidly formed without inputs from grid cells.

Samsonovich and McNaughton (Samsonovich and McNaughton, 1997) proposed a “map-based path integration” model, which associates sensory inputs with a preexisting hippocampal “chart” (a twodimensional attractor map). Although this model and ours share a similar concept, we also provide the biologically plausible mechanism of dendritic computation and inhibition that can enable fast and robust association. Moreover, the chart model has no plastic recurrent connections and hence does not account for replay events. Káli and Dayan (Káli and Dayan, 2000) proposed to train recurrent weights through correlations among DG-to-CA3 inputs, and EC-to-CA3 weights through correlations between DG inputs and EC inputs. Thus, their learning rule also results in correlating EC-to-CA3 inputs with recurrent inputs. However, our learning rule, but not theirs, explains the extremely sparse activity of DG granule cells in spatial exploration (Diamantaki et al., 2016). In our model, DG neurons only occasionally fire to trigger specific firing sequences. This is consistent with the recent finding that a granule cell in DG has at most a single place field (Senzai and Buzsáki, 2017). In addition, our model uses strong recurrent synapses for rapid learning, while their model requires weak recurrent inputs during the early phase of learning.

Several models explain the generation of dendritic calcium spikes (Larkum, 2013; Shai et al., 2015), but the functional implications of calcium spikes were rarely explored. Urbanczik and Senn (Urbanczik and Senn, 2014) proposed a two-compartment model in which dendritic synapses learn to predict somatic activity through unidirectional soma-dendrite interactions. In contrast, our neuron model enables simultaneous learning of somatic and dendritic synapses through bidirectional soma-dendrite interactions. This raises a conceptual difference between the two models: our model performs unsupervised learning of the two input streams, while their model obeys supervised learning of dendritic input using somatic input as a teacher signal. On the other hand, dendritic computation and recurrent networks were combined to improve the capacity of pattern completion (Kaifosh and Losonczy, 2016). However, to our knowledge, the role of dendritic computation in sequence learning was unexplored.

In sum, our multi-compartment learning rule extends the computational ability of neurons to a conjunctive analysis of synaptic inputs targeting different dendritic sites. Because the proximal (somatic) and distal dendrites in pyramidal neurons are targeted by outputs of distinct brain regions, our learning rule has implications for the mechanisms of integrating parallel distributed processes across the brain.

## Methods

### Weight changes and moving thresholds

In all numerical simulations, we modified excitatory synapses in the somatic and dendritic components according to the following second-order stochastic dynamics incorporating delays, weight decays and spontaneous fluctuations:

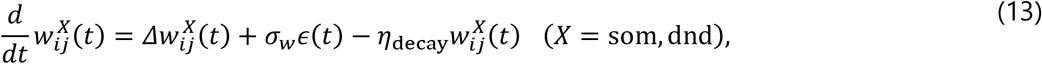

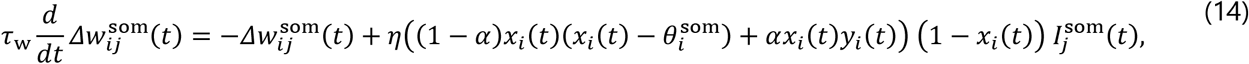

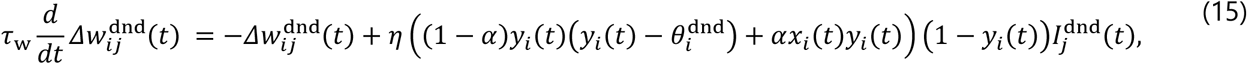

where τ_w_ is the time constant for delays of synaptic changes, η_decay_ is the speed of weight decay, *∊(t)* is normal Gaussian noise and *σ_s_* is the standard deviation of spontaneous fluctuation. The weights were constrained in non-negative values during simulations.

Long-term plasticity of dendritic inhibitory weights 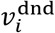 was implemented as

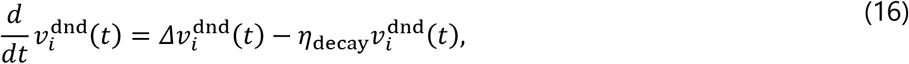

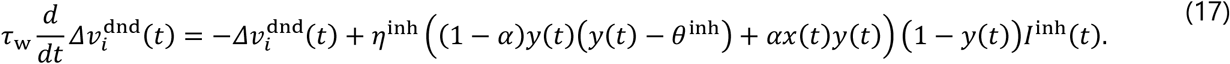

Somatic inhibitory weights 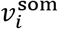 were fixed.

For single-compartment neuron, plasticity follows BCM rule:

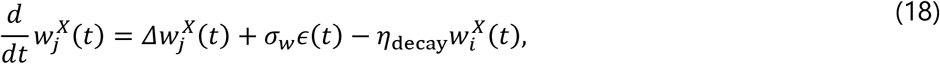

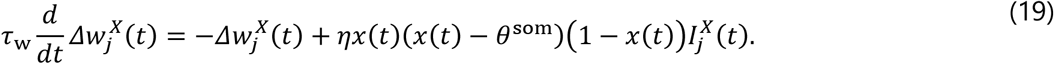

Moving thresholds for BCM theory were defined as 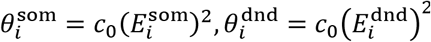. We updated the mean activities 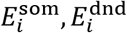 by solving

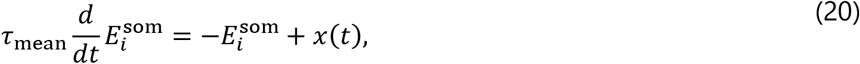

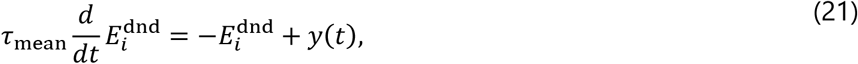

where r_mean_ determines the typical time scale of the averaging.

### Two-compartment recurrent neural network model

Here we define the two-compartment neural network used in Figures 3 to 6. In Figure 1 and Figure 2, we used the model described in the main text without short-term plasticity, NMDA synapses and GABA-B synapses. The activity of neuron *i* in two-compartment recurrent neural networks was described as

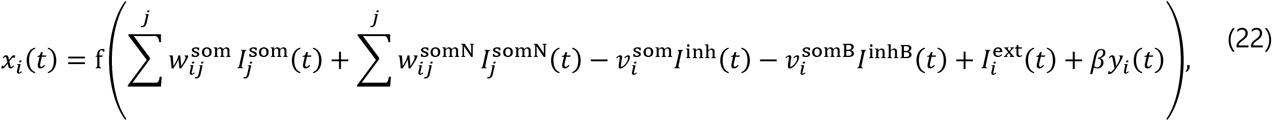

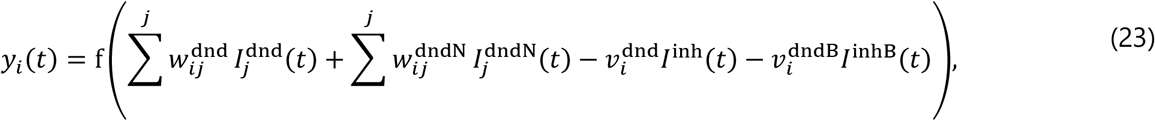

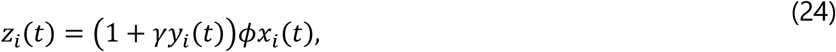

where parameters were set as 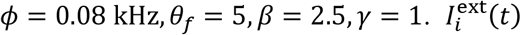 is external input, which varied depending on the simulation settings. Synaptic inputs 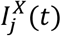 were calculated from recurrent inputs 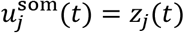 and firing rates of EC neurons 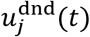 as

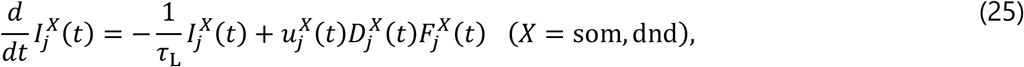

where 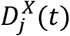 and 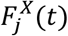 are variables for short-term synaptic plasticity:

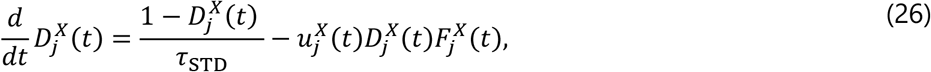

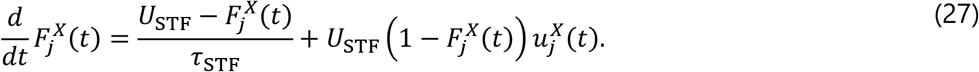

The values of parameters were set as τ_std_ = 500 ms, τ_STF_ = 150 ms and τ_l_ = 10 ms. The value of initial release probability *U*_STF_ was 0.5 for all excitatory synapses in the immobile state of animal, and was changed to 0.03 for recurrent synapses during animal’s movement. At the moment that the animal started a movement from immobile state, 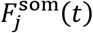 was immediately changed to 0.03. We note that the long-term plasticity rules also depend on short-term plasticity through 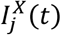.

NMDA currents 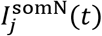 and 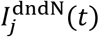 were calculated in a similar way to 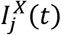 by using a longer time constant 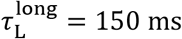. Their weights 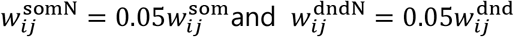 were determined from initial synaptic weights and fixed during all simulations.

Inhibitory feedback *I^inh^(t)* (GABA-A) and *I^inhB^(t)* (GABA-B), were represented by the summation of outputs from the recurrent network as

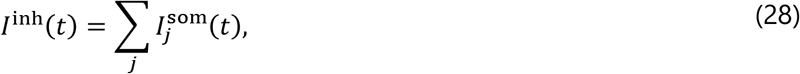

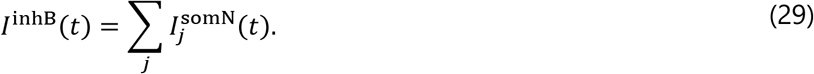

Initial values for 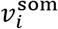 and 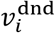 were 4 and 0, respectively, in all simulations. Weights for 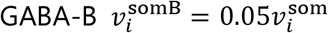 and 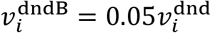 were determined from initial synaptic weights and fixed during all simulations.

The firing rate of EC neurons 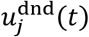x were calculated as

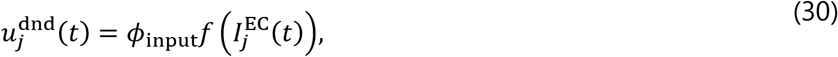

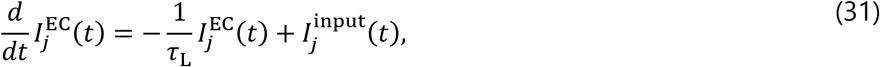

where 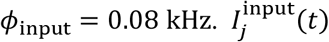 depends on simulation settings.

The values of parameters for plasticity were *α* = 0.9, *r*_0_ = 0.05, *τ*_w_ = 1000 ms, *σ*_w_ = 0.001, *η*_decay_=10^−7^, *θ*_inh_=0.5, *η*_inh_=0.01*η* and *τ*_mean_=60000 ms. Self-connections 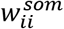 were fixed at zero.

Simulation without dendritic inhibition was performed with η^inh^ = 0.

### Single-compartment recurrent neural network model

In single-compartment recurrent neural networks, all somatic and dendritic inputs were connected to a single compartment (soma). Accordingly, the activity of neuron *i* was described as

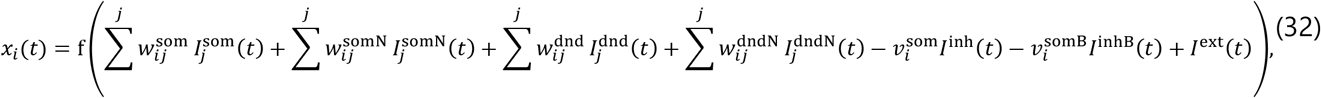

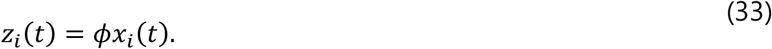

Variables in this model were calculated in the same way to those in the two-compartment model. We updated both somatic and dendritic excitatory weights 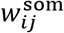 and 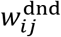 by BCM theory with somatic activity. The values of parameters for the single-compartment model was basically the same as those of the two-compartment model, except ø = 0.09 kHz. Learning speed was set as *η* = 0.5 in Fig. 3D, though different values *η* = 0.1,1.0 were also used in a quantitative assessment.

### Details of the single-cell simulations (Figure 1)

Four independent source signals s_i_(t)(*i* = 1,2,3,4) were generated from Ornstein-Uhlenbeck process

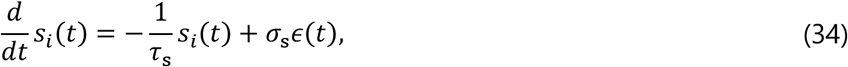

where τ_s_ = 10 ms,σ_s_ = 0.1 and ∊(t) is normal Gaussian noise. Input currents to somatic input neurons 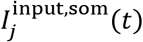 were determined as

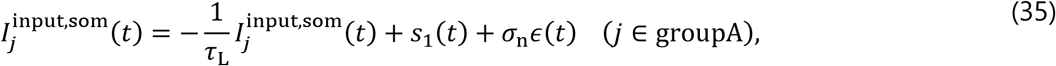

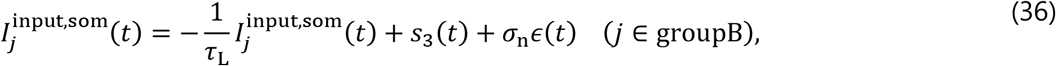

where σ_n_ = 0.1. Input currents to dendritic input neurons 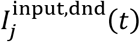 were was determined as

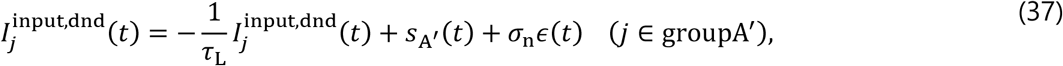

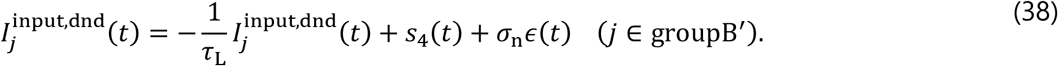

In the case of uncorrelated A and 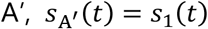 while in the case of correlated A and 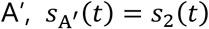. Output firing rates of input neurons 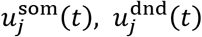 were calculated by the same sigmoidal function f(I) as that of the two-compartment neuron model:

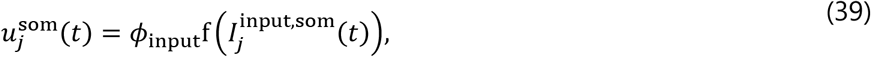

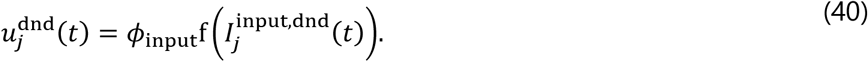

The values of parameters for the two-compartment neuron model were given as *τ*_L_ = 10 ms, *ϕ* = 0.08 kHz, *ɸ*_input_ = 0.08 kHz, *θ_f_* = 5, *η* = 0.2, *γ* = 1, *β* = 0 *α* = 0.5, *r*_0_ = 0.05, *τ*_w_ = 1000 ms, *σ*_w_ = 0.005, *η*_decay_=10^−7^, and *τ*_mean_=60000 ms. Simulations of the single-compartment neuron were performed for *α* = *γ* = 0 without changing the values of the other parameters. Initial weights were uniformly sampled from [0, 5].

### Details of simulations of inhibitory feedback model (Figure 2)

We calculated source signals s_i_(t) in the same way with previous section. In the two-cell simulation for separation, we prepared two source signals. We calculated 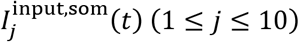 and 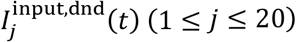 by

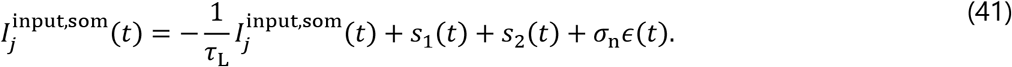

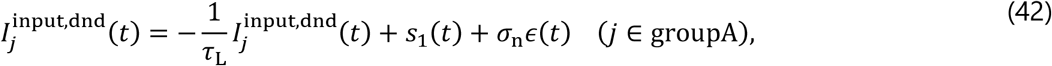

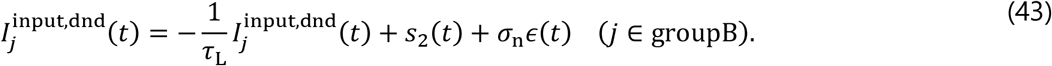

In the single-cell simulation for stabilization, we prepared 21 source signals. Throughout the simulation, we calculated 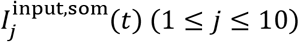 and 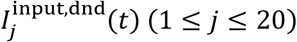 by

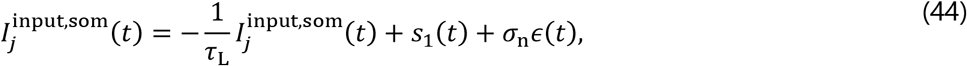

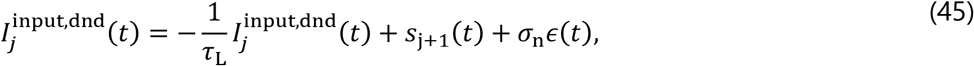

from 0 s to 300 s,

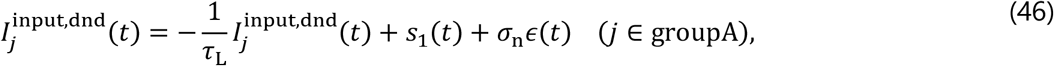

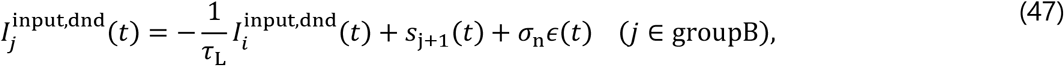

from 300 s to 600 s, and

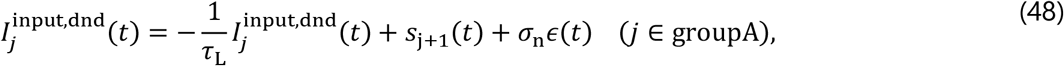

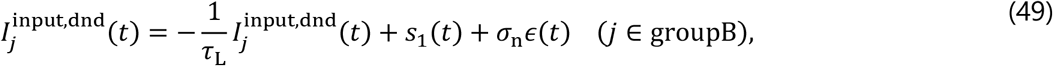

from 600 s to 1200 s. We determined output firing rates of input neurons 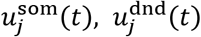 by Eq. (39) (40).

The values of parameters for the two-compartment neuron model were given as 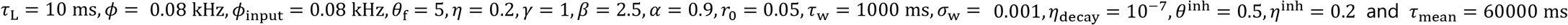. Initial excitatory weights were uniformly sampled from [0, 5] and initial dendritic inhibitory weights 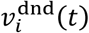 were zero. Somatic inhibitory weights 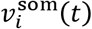 were fixed to 20. Simulation without dendritic inhibition was performed with η^inh^ = 0.

### Simulation settings for the one-dimensional track (Figure 3)

We used 300 CA3 neurons and 500 EC neurons. Initial recurrent synaptic weights from neuron *j* to neuron *i (i ≠ 7)* in CA3 were given as

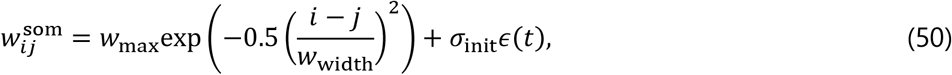

where w_max_ = 18, w_width_ = 5 and σ_init_ = 0.05. Here we included random fluctuation of weights sampled from normal Gaussian distribution ∊(t), and negative weights were set to zero. Self-connections 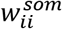 were always zero. In qualitative assessment, we multiplied 0, 0.5, 0.75, or 1.25 to w_max_ in each simulation.

Initial synaptic weights from EC 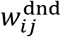 were firstly determined as

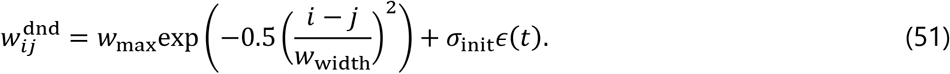

We used this setting to simulate the "familiar track". In the simulation of "unfamiliar track", we randomly shuffled values of these weights in each postsynaptic neuron *i*. Namely, shuffling was performed for index y.

A function satisfying 0 ≤ *pos(t)* ≤ 1 designated the animal’s position on the one-dimensional track. The animal stopped at *pos(t) = 0* from 0 s to 5 s. From 10s to 25 s, the position in first run is expressed as

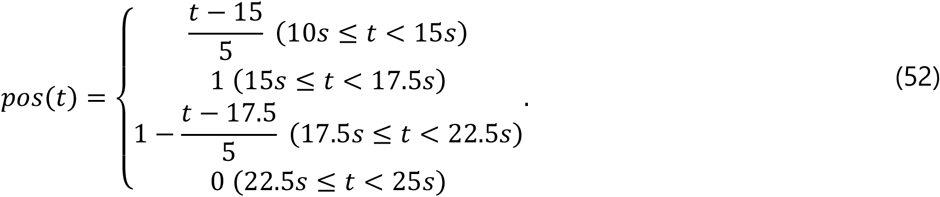

From 25s to 40 s, the position in second run is expressed as

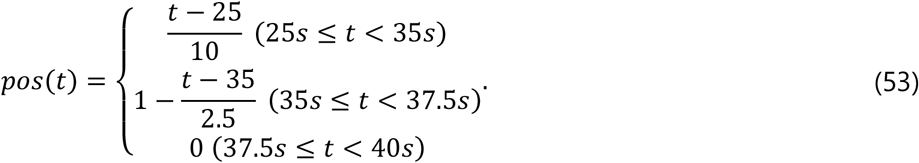

From 40s to 50 s, the position in third run is expressed as

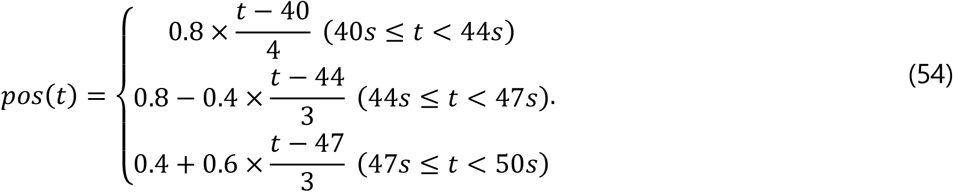

External inputs to somatic compartments 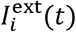 were

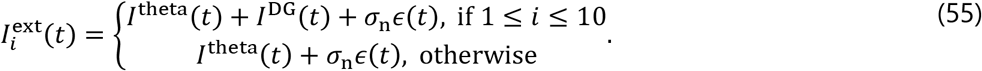

*I*^DG^(*t*) is input from dentate gyrus (DG), which takes 0.5 when DG is active and 0 otherwise. When the animal was running in the portion *pos(t)* < 0.05, DG was continuously activated. When the animal was stopping in *pos(t)* < 0.05, the activation of DG follows a Poisson process at 1 Hz, and the duration of each activation was 10ms.

*I*^theta^(*t*) stands for theta oscillatory input from medial septum

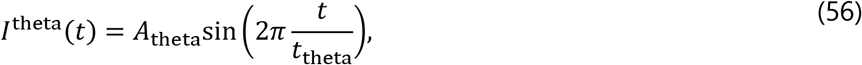

during run and *I*^theta^(*t*) = 0 during immobility.

Inputs to EC neurons 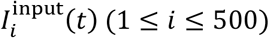 were given as

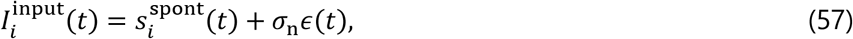

during immobility. During run, Inputs to position-dependent EC neurons 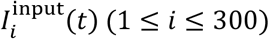 were given as

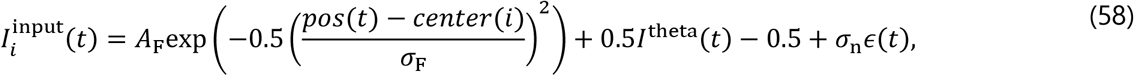

where ∊(t) is normal Gaussian noise and *center* 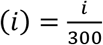. Inputs to distractor EC neurons (301 ≤ *i* ≤ 500) during run were given as

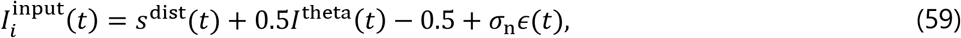

Sources for spontaneous activity 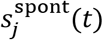 and distractors *s*^dist^(*t*) were generated from independent Ornstein-Uhlenbeck processes

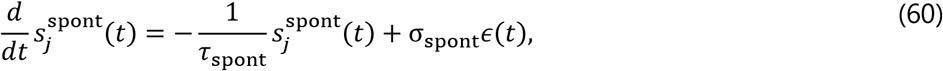

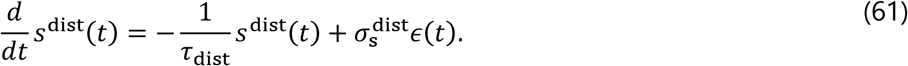

Parameters were 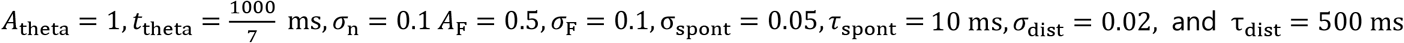

### Simulation settings for spontaneous replay (Figure 4)

Initial setting was the same as in Fig. 3. In Fig. 4, run was finished after the first run on the one-dimensional track, and spontaneous activity was simulated for the next 600 s. After that, simulation of the “third run” in Fig. 3 was conducted. During spontaneous activity, simulation setting was basically the same as that of immobility periods in Fig. 3. However, we added population bursts in EC, which were simulated by transiently changing *σ*_spont_ from 0.05 to 0.1. The occurrence of population bursts followed a Poisson process at 1 Hz, and each burst lasted for 200 ms. For the results shown in Fig. 4D, all weights of the dendritic inhibition and *η*^inh^ was set to zero when 60 s elapsed after the beginning of spontaneous activity, and simulation of the “third run” in Fig. 3 was conducted afer that.

### Simulation settings for the Y-shape track (Figure 5)

We used 450 CA3 neurons. We divided these neurons into three groups, 1 ≤ *i* ≤ 150,151 ≤ *i* ≤ 300,301 ≤ i ≤ 450, and recurrent synaptic weights within each group were determined in the same way as in the onedimensional track, using *w*_max_ = 18, *w*_width_ = 5 and *σ*_lnit_ = 0. Recurrent synaptic weights across groups were initially zero. Initial synaptic weights from 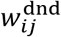 were uniformly sampled from the interval [0, 2].

The current position of the animal on the Y-shape track was specified by the arm number (*arm (t)* = 1,2,3) and the position on the current arm, 0 ≤ *pos(t)* ≤ 0.5. During the first 10 s, the animal stopped at the center of the Y-shape arm. After that, the animal repeated the following movement on different arms every 10 s:

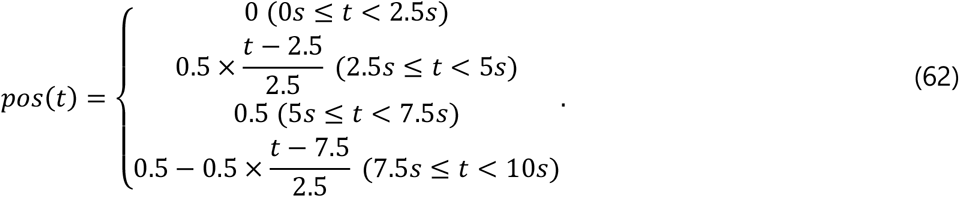

External inputs to the somatic compartments 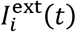 were basically the same as the ones used for the onedimensional track except that DG inputs with strength 1 were induced to 10 neurons per group, 1 ≤ *i* ≤ 10,151 ≤ i ≤ 160,301 ≤ *i* ≤ 310 in the region 0 ≤ *pos*(t) ≤ 0.05 on all arms.

We used 450 position-dependent EC neurons, and inputs to these EC neurons 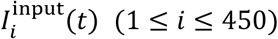 during immobility were the same as those in the one-dimensional track, whereas the inputs during the run depended on animal’s position as

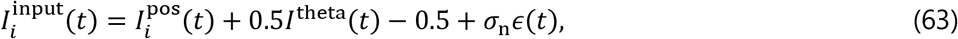

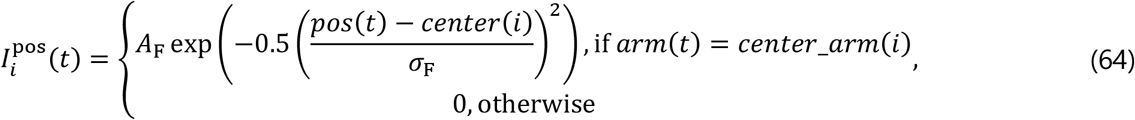

where the receptive field center *center(i)* of neuron *i* was uniformly sampled from [0, 0.5] and 150 neurons were assigned to each arm: *center_arm(i)* = 1, 2 and 3 for 1 ≤ *i* ≤ 150, 151 ≤ *i* ≤ 300, and 301 ≤ *i* ≤ 450, respectively.

### Simulation settings for branching firing sequences (Figure 6)

We used 400 CA3 neurons and 300 EC neurons. We divided neurons into three groups, 1 < *i* < 100 (root), 101 ≤ i ≤ 250 (branch A), 251 ≤ *i* ≤ 400 (branch B), and recurrent synaptic weights within each group were determined in the same way with the one-dimensional track, using *w*_max_ = 20, *w*_width_ = 5 and σ_lnit_ = 0. Weights were set to zero between branch A and branch B, and

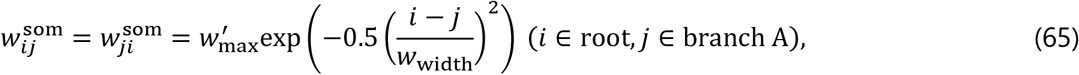

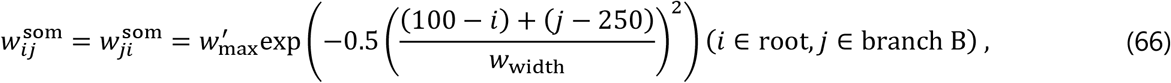

for other weights. The value of 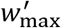 was 18. Initial synaptic weights from EC 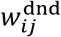 were uniformly sampled from the interval [0, 2].

The animal was immobile from 0 s to 60 s, and from 75 s to 135 s. In these periods, the number of firing sequences in each branch was counted to compare the propagation of firing sequences before and after an experience. During 60 s – 75 s (first experience) and 135 s – 150 s (second experience), **pos*(t)* was changed in a same way with “first run” on the one-dimensional track (Fig. 3). *center(i)* for each neuron *i* was uniformly sampled from [0, 1] for the first experience, and resampled in a similar way for the second experience. Other simulation settings were basically the same as in simulations of the one-dimensional track.

### Information per spike

We evaluated the accuracy of place fields by using information per spike given as follows:

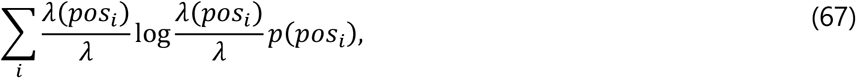

where *pos_i_* is the binned position of the animal (*i = 1, …,N_bin_*), *p(pos_i_)* is the probability that the animal is found at given position *i*, λ is the mean firing rate of the cell, λ(pos_i_) is the mean firing rate when the animal is in *pos_i_*. After removing immobile periods, we computed information per spike for all CA3 neurons having the mean firing rate higher than 1 Hz and averaged this quantity over these neurons. The number of bins *N_bin_* was 50 in Figure 3 and 75 (25 for one arm) in Figure 5.

### Homeostasis in BCM theory

BCM theory used in this study is slightly different from the conventional one. Therefore, we analyze the homeostasis of our BCM theory:

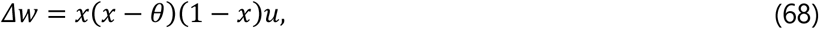

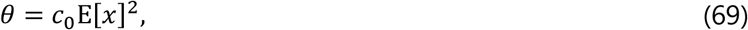

where *x* is postsynaptic activity and *u* is presynaptic input.

As shown in Results, this learning rule is given as a gradient ascent of the objective function:

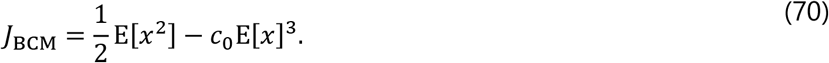

Because *0 ≤ x ≤ 1* in this paper (*x* is given by a sigmoidal function), we use an approximation *E[x^2^] ≈ E[x]* in the above equation. With this approximation, the objective function becomes

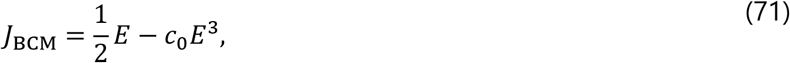

defining *E = E[x]*. The fixed point of this objective function is derived as

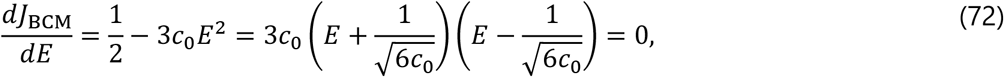

Therefore, we should set 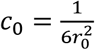 to make the mean firing rate converge to *r_0_*.

If we include the CCA term, the objective function is

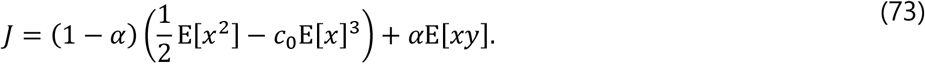

If *x* and *y* are independent, *E[xy] = E[x]E[y]* holds. Assuming a similar learning rule for y, we can approximate this term with *E[x]*^2^, and in this case the objective function becomes

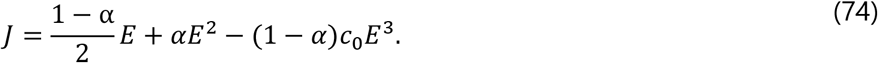

By solving

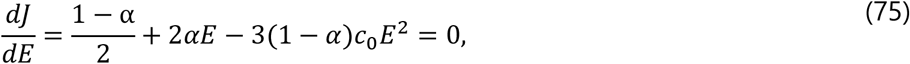

We obtain

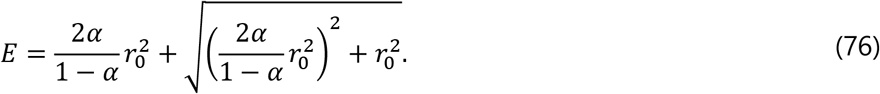

The value of *E* is always larger than *r_0_*, but the difference is moderate if *r_0_ ≪ 1*. For instance, when *r_0_ = 0.05* and *α* = 0.9 (these values were used in this study unless otherwise stated), this value is about 0.11.

If *x* and *y* are perfectly correlated and *x ≈ y*, we can use an approximation, E[*xy*] ≈ *E*[*x*^2^]. Then, the analysis is similar to the previous case without the CCA term if we change the coefficient of E[*x*^2^] in *J*_BCM_ from 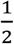 to 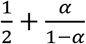. Then, the approximate fixed point changes to 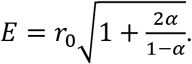. As 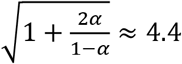 for *α* = 0.9, the equation *E[x] ≈ 4.4r_0_* has a solution only when *4.4r_0_* ≤ 1.

Because the above derivation holds only approximately, the convergence to a stable solution is not really ensured. However, the homeostasis was preserved in the present simulations, and the mean firing rate always converged to a value slightly higher than the theoretical estimation. Note that plasticity of inhibitory synapses is not taken into account in this analysis, and inhibitory plasticity sometimes caused instability for some parameter values.

### Code availability

All codes for simulations and visualization were written in Python 3 and available at https://github.com/TatsuyaHaga/preplaymodel_codes.

## Acknowledgements

We are grateful to K. Inokuchi, S. Fujisawa, and T. McHugh for fruitful discussions, and C. Yokoyama for helpful comments on the manuscript. This work was partly supported by CREST, JST (JPMJCR13W1) and Grants-in-Aid for Scientific Research (KAKENHI) from MEXT (No. 15H04265 and 16H01289).

## Author contributions

T. H. and T. F. equally contributed to the conception and design of the model and the preparation of the manuscript. T. H. performed mathematical calculations, numerical simulations and data analysis.

